# The genetic architecture of trabecular bone score and association with fracture: a genome-wide association and meta-analysis

**DOI:** 10.1101/2025.10.01.678241

**Authors:** Haojie Lu, Griffin Tibbitts, Gloria Hoi-Yee LI, Monika Frysz, Christine W. Lary, Katerina Trajanoska, Nilabhra R. Das, Lissette de Groot, Ching-Lung Cheung, Natasja M. van Schoor, André G. Uitterlinden, Nathalie van der Velde, Jonathan H Tobias, Fernando Rivadeneira, David M. Evans, Douglas P. Kiel, John P. Kemp, Carolina Medina-Gomez

## Abstract

**Background:** Trabecular bone score (TBS) is a texture-based measurement derived from DXA scans, which describes the distribution of mineral across the vertebral bodies. Identifying its genetic determinants is crucial for enhancing understanding of its biological basis and clarifying its relationship with fracture risk.

**Methods:** We conducted a large-scale genome-wide association study (GWAS) of TBS from 44,767 participants (95.08% European ancestry). Sex heterogeneity was assessed through sex-stratified GWAS. Post-GWAS analyses included functional annotation, gene-set enrichment analysis, and *cis*-expression quantitative trait locus (*cis-*eQTL) colocalization. Two-sample and multivariable Mendelian randomization (MR) were employed to investigate the causal effect of TBS on fractures.

**Findings:** We identified 33 independent variants associated with TBS, which explained 4.06% of TBS phenotypic variance. All the discovered signals map to previously identified BMD-associated loci. Additionally, we identified genetic variants in the *RAB11FIP3* locus that exhibited a sex-heterogeneous effect, being associated with TBS exclusively in males. Functional annotation and colocalization identified functional genes related to TBS. The two-sample and multivariable MR analyses indicated that TBS potentially has an independent causal effect on fracture risk.

**Interpretation:** Our study unveiled 33 independent loci associated with TBS, all in BMD-associated loci. One locus was further identified as associated with TBS only in males. MR results suggested that genetically derived TBS may be causally associated with fracture risk at different sites, potentially beyond BMD. This study provides insights into the TBS genetic architecture and uncovers its potential clinical applications in fracture risk prediction.

**Funding:** All funding information can be found in the Acknowledgements section.

**Research in context:** *Evidence before this study:* Osteoporosis is a common disease prevalent in older adults, mainly diagnosed by low bone mineral density and disruption of bone architecture. While the genetic determinants of bone mass have been well studied, the genetic architecture of bone features beyond BMD remains largely unknown.

*Added value of this study:* This study presents a large genome-wide association meta-analysis of the trabecular bone score (TBS), a measurement of trabecular distribution of bone mineralization across vertebral bodies. Our study identified 33 independent variants associated with TBS and one association signal only in males. Functional annotation highlighted crucial genes related to TBS. The two-sample and multivariable Mendelian randomization (MR) analyses suggested that TBS may be causally linked with fracture risk, potentially independent of BMD.

*Implications of all the available evidence:* Our study represents an expansion of the traditional bone density phenotype used in most previous skeletal genetic studies. Based on the findings that all of the significant loci for TBS have been previously identified in GWAS of BMD, we conclude that TBS-associated variants have pleiotropic effects affecting not only mineral density but also the distribution patterns of minerals in the cancellous bone. Continuing to perform genetic studies of new skeletal phenotypes beyond BMD could ultimately impact the management and prediction of osteoporosis patients.

## Introduction

Osteoporosis is a common disease characterized by low bone mass and disruption of bone architecture, which can result in compromised bone strength and an increased fracture risk in the elderly. Bone mineral density (BMD), assessed using dual-energy X-ray absorptiometry (DXA), is employed in clinical practice to diagnose osteoporosis. The genetic determinants of BMD across various skeletal sites have been largely explored in previous genome-wide association study (GWAS)^1–5^.

While BMD is a crucial metric for defining bone health, it does not capture all bone features related to bone strength. Many patients experiencing fragility fractures exhibit normal or osteopenic BMD levels^6^. The assessment of bone quality is complex and still in development, but includes bone architecture, bone turnover, and bone material features^7^. From the imaging perspective, low-cost modalities that intend to reuse conventional DXA images to assess bone features other than BMD, have been proposed^7^. One of these modalities is the trabecular bone score (TBS), a gray-level textural measurement derived from lumbar spine (LS) DXA scans^8,9^.

TBS is an FDA-approved software add-on marketed by TBS iNsight (Medimaps, Plan-les-Quates, Switzerland). It provides information on the heterogeneity and distribution of bone mineral within a vertebral body, and can differentiate bone structures that exhibit the same areal BMD. In clinical studies, TBS has been shown to be associated with fracture risk and predict risk of fracture over traditional fracture risk assessment tools (FRAX) and areal BMD^10,11^.

In this study, we conducted the most extensive TBS GWAS to date, involving 44,767 individuals of predominantly European background across multiple cohorts. We also performed sex-stratified GWAS to identify variants exhibiting sex-heterogeneity. Subsequently, we completed in-silico analyses to (1) elucidate the biological pathways through which genes in identified genetic loci might be acting, and (2) assess the causal effect of TBS on fracture risk.

## Methods

### Study populations and TBS measurement

This study involved participants from eight population-based studies, each with comprehensive data on TBS, relevant covariates (age, sex, and study-level covariates), and genetic data. These studies included the Rotterdam Study (RS) I, II, and III, Generation R-mothers (GenR), Framingham Heart Study (FHS), B-Vitamins for the PRevention Of Osteoporotic Fractures(B-PROOF), UK Biobank (UKBB), and Hong Kong Osteoporosis Study (HKOS). Altogether, data from 42,564 individuals of European ancestry (23,446 females) and 2,203 individuals of non-European ancestry (1,736 females) were analyzed. Genotyping of the participants was completed on different platforms and imputed using different reference panels, including 1000 Genomes Project (1KGP, Phase 3, version 5) and the HRC release 1.1.

TBS is derived from LS-BMD scans using the TBS software (iNsight; Medimaps Group, Geneva, Switzerland), which provides information on the heterogeneity and distribution of bone mineral within a vertebral body. The TBS calculation adjusts for soft tissue effects through two main approaches: utilizing BMI as an indirect measure of soft tissue thickness (TBS_BMI_) or directly incorporating soft tissue thickness (TBS_thickness_)^12^. Information on phenotype acquisition per cohort is provided in Supplementary Table 1.

### Genome-wide association studies and meta-analyses

We used additive models to test the association between genetic variants and TBS. For each cohort, TBS measurements were standardized to have a mean of 0 and a variance of 1 across all individuals (i.e., males and females together); and males and females separately, for the sex-stratified analyses. All our models were adjusted for sex (when pertinent), age, age squared, genetic principal components, and study-level covariates (Supplementary Table 1). We employed the R package EasyQC (version 23.8) to conduct quality control on summary statistics of each study prior to the meta-analysis^13^. Single nucleotide polymorphisms (SNPs) with minor allele frequency (MAF) below 0.5% or imputation quality score less than 0.4 (Impute) or less than 0.3 (Minimac) were excluded. Moreover, SNPs with an absolute value of estimated effects or standard errors greater than 10 were excluded from the meta-analysis.

The individual cohort GWAS summary results were meta-analyzed using an inverse-variance weighted model implemented in METAL (version 2020-05-05). We also conducted a European-specific meta-analysis (N = 42,564) using only summary statistics from studies comprising individuals of European ancestry, which served as the backbone for subsequent investigations. In this last group, we additionally performed a sex-specific GWAS meta-analysis (N_male_ = 19,118; N_female_ = 23,446). For the final quality control of results from meta-analysis, we excluded insertion-deletion variants, variants with MAF less than 0.5%, showing heterogeneity across multiple cohorts (I^2^ >30), with a sample size less than 4,477 (i.e., 10% of the total individuals) or present in only one cohort. Variants with an association P-value less than 5×10^−8^ were considered genome-wide significant (GWS).

Next, we used the linkage disequilibrium (LD) score regression implemented in LDSC^14^ (version 1.0.1) to quantify the genomic inflation contributed by population stratification, cryptic relatedness, and other potential factors. We also estimated the SNP-based heritability (h^2^-SNP) of TBS, i.e., the proportion of phenotypic variance explained by the variants included in the GWAS meta-analysis. The analysis was conducted using pre-computed LD scores derived from the European samples in the 1KGP. All analyses were limited to high-quality variants present in the HapMap3 Project^15^, outside the major histocompatibility complex (MHC) region on chromosome 6 (6p21.3).

### Conditional and joint analysis

We used conditional and joint association analysis (COJO) from GCTA (v1.94.1) to identify conditionally independent variants associated with TBS in our European-based meta-analysis^16,17^. The LD-reference was built based on 378,485 unrelated European individuals (KING kinship coefficient <0.044) from the UKBB (project No. 17295). Variants were excluded from the reference panel based on Hardy-Weinberg proportions (P-value < 1×10^−6^), missing call rate (< 95%), or MAF (< 0.01%, with a minor allele count > 1). A strict QC was then performed using DENTIST (v1.2) to compare the variants from the meta-analysis summary statistics and the generated LD reference^18^. Genetic variants were excluded from the COJO analysis if their alleles did not match, allele frequencies differed by more than 10%, or were predicted to be incorrectly imputed. Parameters in COJO analysis included a window size of 10,000 kb and multiple regression collinearity of 0.8. The TBS phenotypic variance explained by each independent associated SNP was estimated using the following formula^19^ (1):

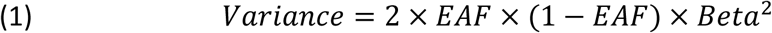

where *EAF* and *Beta* represent the effect allele frequency and estimated effect size of each variant. As TBS was standardized (with phenotypic variance of 1), the total proportion of variance explained by GWS-independent SNP corresponds to the summation of (1) across all the lead variants.

To investigate if TBS-variants are independent of known BMD-variants, we also conducted an additional COJO analysis using the same strategy but conditioning on independent GWS variants from the well-powered GWAS of estimated heel BMD (e-BMD) in UKBB^20^. The remaining GWS variants were then checked against the publicly available results of DXA-derived BMD GWAS^2–4^.

### Sex heterogeneity

In addition to conducting sex-stratified GWAS meta-analyses, we used Easy-strata^21^ to identify whether the sex-specific GWS loci presented sex heterogeneity, using equation (2):

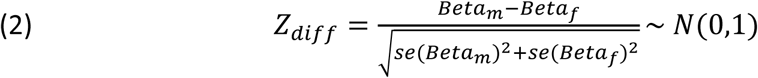

Where *Beta*_*m*_and *Beta*_*f*_ are the estimated effect sizes for the SNPs in males and females GWAS, and *se*(*Beta*_*m*_) and *se*(*Beta*_*f*_) are the standard errors for the estimated effect sizes. To account for multiple comparisons, we applied Bonferroni correction considering the number of sex-specific GWS loci tested (n = 22; significance threshold P = 2.27 × 10⁻³) for identifying sex-heterogeneous loci.

### Functional annotation

The Functional Mapping and Annotation of GWAS (FUMA) (version v1.5.2) was used for the functional annotation of TBS-associated variants^22^. We first applied the “SNP2GENE” function in FUMA to identify the GWS loci from multi-ancestry and sex-stratified TBS GWAS, with default parameter settings and a window of 10,000 kb. The 1KGP was used as an LD reference panel in FUMA, including 1KGP-ALL (for multi-ancestry) and 1KGP-EUR (sex-specific).

Through the FUMA pipeline, all significant variants were annotated with ANNOVAR^23^ to predict their functional consequences (e.g., affecting the coding regions or the splicing of a protein). Also, we annotated each lead variant with the Combined Annotation Dependent Depletion (CADD) score, a framework that assesses the deleteriousness of genomic variants (variants with CADD scores larger than 12.37 are considered highly deleterious)^24^. As most of the GWS variants lie in DNA non-coding regions, we also used the public database (https://regulome.stanford.edu/regulome-search/) of RegulomeDB (version 2.2) to predict their potential functional annotation^25^.

To test whether the TBS loci-mapped genes were enriched in particular biological pathways or functional categories, we performed the gene-set enrichment analysis using GENE2FUNC function in FUMA^22^. The gene sets used were obtained from public databases, such as MSigDB and WikiPathways^22^.

### eQTL and Colocalization analyses

To determine whether TBS-associated SNPs influence the expression of neighboring genes in different tissues (driven by shared causal variants), we performed colocalization with expression quantitative trait locus (eQTL) information from GTEx^26^ (version 8). We focused on the eQTL datasets in whole blood (N = 573, femoral vein) and skeletal muscle (N = 602, gastrocnemius muscle), considering their functional and biological relevance with skeletal traits. In addition, we also queried eQTL data from osteoclast-like cell cultures differentiated from peripheral blood mononuclear cells obtained from 158 women^27^. We only focused on cis-eQTLs, i.e., defined by a window size of 1,000kb around each conditionally independent lead variant.

The colocalization was performed using Pair-Wise Conditional analysis and Colocalisation analysis (PWCoCo), which combines conditional and colocalization analyses within a Bayesian framework, allowing the analysis of non-primary signals^28^. We used the default prior settings: P_1_ = 1.0 × 10^−4^, P_2_ = 1.0 × 10^−4,^ and P_12_ = 1.0 × 10^−5^, where P_1_, P_2_, and P_12_ represent the prior probability of a variant being causally associated with TBS, tissue-specific gene expression, or both traits separately. Colocalization analyses leveraged variants from the TBS GWAS and cis-eQTL of the different tissues described above. It tests five hypotheses, i.e., no causal signal associated with any of the traits (H0), one causal signal associated with one specific trait (H1 and H2), different causal signals associated with both traits (H3), and one shared causal signal associated with both traits (H4). A posterior probability (PPH4) larger than 0.8 is considered strong evidence of colocalization.

For these analyses, an LD reference panel was constructed using imputed (HRC 1.1) data from participants with a North European background in the Rotterdam Study I (RS-I). We defined the North-European background by calculating the mean and SD of the CEU samples present in HapMap across the first four principal components. Any RS-I participant deviating more than 4 SD from the mean of this population in any of these four principal components was excluded from the reference panel. Next, we calculated the pairwise Identity-by-descent (IBD) distance between participants and randomly excluded one individual from every pair with an IBD proportion larger than 0.35. A total of 5,775 participants from RS-I were finally kept. In addition, we filtered variants with low imputation quality (rsq < 0.3).

### Genetic correlation

In a subset of UKBB participants (N = 28,534), we first estimated the genetic correlation (rg) of TBS measured using different adjustments: TBS_BMI_ and TBS_thickness_. We then estimated the pairwise rg of TBS (both sex-combined and sex-stratified) with BMD from different skeletal sites, including LS-BMD^4^, FN-BMD^4^, e-BMD^20^, total body (TB-) BMD^2^ and skull-BMD^3^, as well as fractures that happened at any bone (AnyFX)^1^, hip bone (HipFX)^29^, and forearm bone (ForearmFX)^30^. Other traits surveyed included body height^31^, body mass index (BMI)^31^, whole body lean mass^32^, appendicular lean mass^32^, hand grip^33^, 25-hydroxyvitamin D concentration (VitD)^34^, type 2 diabetes (T2D)^35^, coronary artery disease (CAD)^36^ and coronary artery calcification (CAC)^37^. Additionally rg between those phenotypes and LS-BMD, FN-BMD, and e-BMD were calculated. A Bonferroni multiple-testing correction considering 17 traits (P-value < 2.94×10^−3^) was used as a significance threshold. These analyses were carried out using LD Score regression^38^.

### Mendelian randomization investigating the causal effects of TBS on fractures

We used two-sample MR to investigate the potential univariate causal effect of TBS on fracture risk from different bone sites, including AnyFX^1^, HipFX^29^, and ForearmFX^30^. Similar analyses were run for FN-BMD^4^. We restricted the analysis to SNPs available from the corresponding summary statistics of TBS and FN-BMD. The lead GWS SNPs at each locus (with a window size of 10,000kb) were included as genetic instrumental variables (IVs) for the two-sample MR. We first harmonized IVs effect alleles with conflicting directions (e.g., T/G for TBS or FN-BMD and G/T for fracture) or strand issues (G/T for TBS or FN-BMD and C/A for fracture) to make sure the estimated effect size from both exposures and outcomes were referring to the same effect alleles^39,40^. Additionally, we removed palindromic variants with intermediate allele frequency. To guard against reverse causation, we used the MR Steiger test to filter variants that explained more variance in fractures than in the exposures^41^. The MR-PRESSO outlier test with a strict threshold (P-value less than 1) was used to identify and remove outlier IVs with a heterogeneity effect^42,43^.

We employed the inverse variance-weighted (IVW) method to combine Wald estimates (i.e., the SNP-fracture regression coefficient divided by the SNP-exposure regression coefficient)^44^ for each IVs and estimate the overall causal effect of TBS on different fracture risk.

We used additional sensitivity analyses to assess the validity of the included IVs. Cochran’s Q-statistics based on the IVW and MR-Egger regression were used to test the IVs’ heterogeneity^45,46^. The IVs pleiotropic effects were tested using the intercept from MR-Egger regression, as a nonzero intercept suggests the presence of unbalanced horizontal pleiotropy^47^. To test whether there was a weak instrument bias in the two-sample MR, we calculated the F-statistic^41^ using the formula (3):

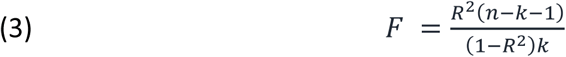

Where *n* and *k* are the sample size and the number of IVs included in MR; *R*^2^ is the TBS variance explained by the IVs and calculated from formula (1). An F-statistic larger than 10 indicates that substantial weak instrument bias is unlikely^41^.

The Python-based package GWASLab (version 3.4.24) and the R packages TwoSampleMR (version 0.5.6) and MRPRESSO (version 1.0) were used to conduct the described analysis.

### Multivariable Mendelian randomization

Using the same set of GWS SNPs as IVs, we applied multivariable Mendelian randomization (MVMR)^48^ to further investigate the direct causal effect of TBS on AnyFX^38^, HipFX^29^, and ForearmFX^30^ independent of BMD, by including FN-BMD^4^ as co-exposure. As some lead SNPs from TBS or FN-BMD mapped closely, we clumped the instruments (LD r^2^ of 0.01 and a 10,000kb window) using the 1KGP European individuals as the reference panel. The clumping was performed based on the TBS association strengh. Similarly, the strict threshold (P-value < 1) from the outlier test in MR-PRESSO (version 1.0) was applied to detect and remove IVs with heterogeneous effects. We performed a second round of clumping using the FN-BMD reported associations as a sensitivity analysis.

We utilized the inverse-variance weighted method as the primary model for MVMR. The conditional F-statistic was used to evaluate the weak IVs bias, which represents the independent proportion of variance explained in each exposure after accounting for associations with the other exposures^49^. An F-statistic larger than 10 indicates less evidence of weak instrument bias. Due to the high genetic correlation between TBS and FN-BMD, we also applied the MVMR-Q(het) method as a sensitivity analysis. This method provides a reliable causal effect in the presence of weak instruments and heterogeneity^48^. This method estimates the causal effect by minimizing the heterogeneity Q-statistic from the input of IVs association with exposures and outcome^48^. We estimated the covariance between exposures using the phenocov_mvmr() function from R package of MVMR, with a phenotypic correlation of 0.62 between TBS and FN-BMD calculated from RS (N = 6,719).

Finally, we also performed a sensitivity analysis using e-BMD as a coexposure in MVMR, rather than FN-BMD, since e-BMD represents the most large-scale GWAS of BMD to date. Lead variants from each locus associated with e-BMD (based on the original research significance definition of P-value < 6.6×10^−9^) were included in MVMR^20^. Similarly, we estimated the covariance between exposures based on the phenotypic correlation of 0.36 between TBS and e-BMD calculated from UKBB (N = 28,182).

The Python-based package GWASLab (version 3.4.24) and the R packages of TwoSampleMR (version 0.5.6), MendelianRandomization (version 0.10.0), MVMR (version 0.4), and MRPRESSO (version 1.0) were used to execute the described analyses.

### Ethics

Written informed consent was obtained from all participants and all study protocols were approved by the institutional review boards of each centre. Further details are available in the Supplementary Materials.

## Results

### GWAS and meta-analysis

The high genetic correlation (rg: 0.999, P-value < 1.79 × 10^−308^) between TBS measurements from different software versions (adjusted for BMI or tissue thickness) in UKBB (N = 28,534), supported the combination of all available data, regardless of the TBS software version used (Supplementary Figure 1). After meta-analysis and quality control, we analyzed genotypes of 44,767 individuals and 8,535,982 variants. Detailed information on each cohort is provided in Supplementary Table 1 and Supplementary Methods. The description of the participants’ characteristics can be found in Supplementary Table 2.

We identify 29 loci (Supplementary Table 3 and Figure 1) associated with TBS in the multi-ancestry meta-analysis, with no evidence of genomic inflation (λ_GC_: 1.04). In the European subset (N = 42,564). The COJO analysis further identifies 33 conditionally independent SNPs associated with TBS, altogether explaining 4.06% of the TBS phenotypic variance (Table 1). The TBS SNP-based heritability was estimated as 20.01% (95% CI: 15.97% to 24.13%), whereas the LD-score regression intercept was 0.97 (SE: 0.01), suggesting unlikely inflation due to population stratification, cryptic relatedness, or latent confounding.

**Figure 1:**
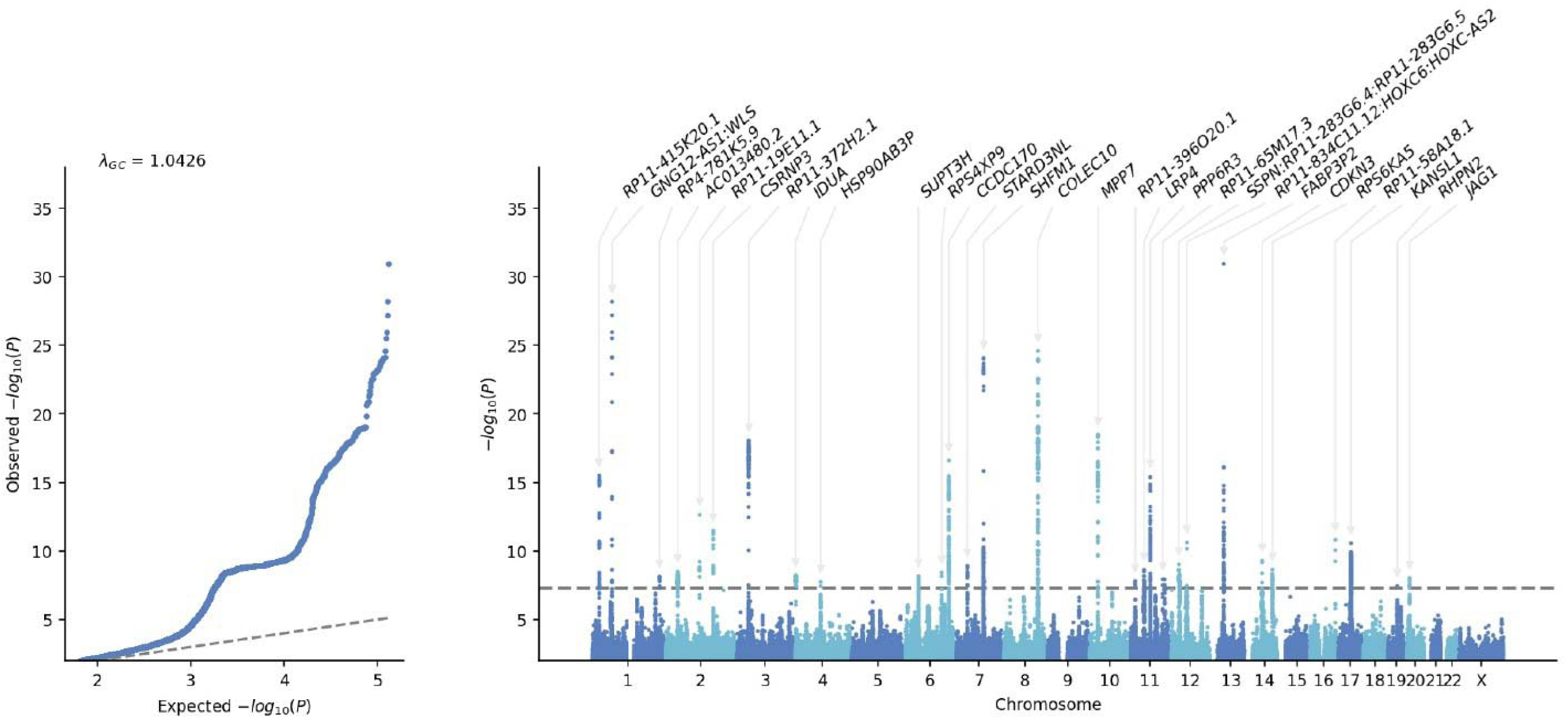
Quantile-quantile plot (left) and Manhattan plot (right) of multi-ancestry TBS genome-wide association study from multi-ancestry (N = 44,767). In Quantile-quantile plot (left), y-axis refers to the -log10 transformed P-value from TBS GWAS, x-axis represents the -log10 transformed expected P-value (two-sided). In the Manhattan plot, the y-axis corresponds to the - log10 transformed P-value of each SNP association, and the x-axis corresponds to the base-pair position along the chromosome. The dashed line indicates genome-wide significance (P-value < 5×10-8). The genes in the Manhattan plot refer to the closest gene to the lead variants of each locus.

**Table 1.**
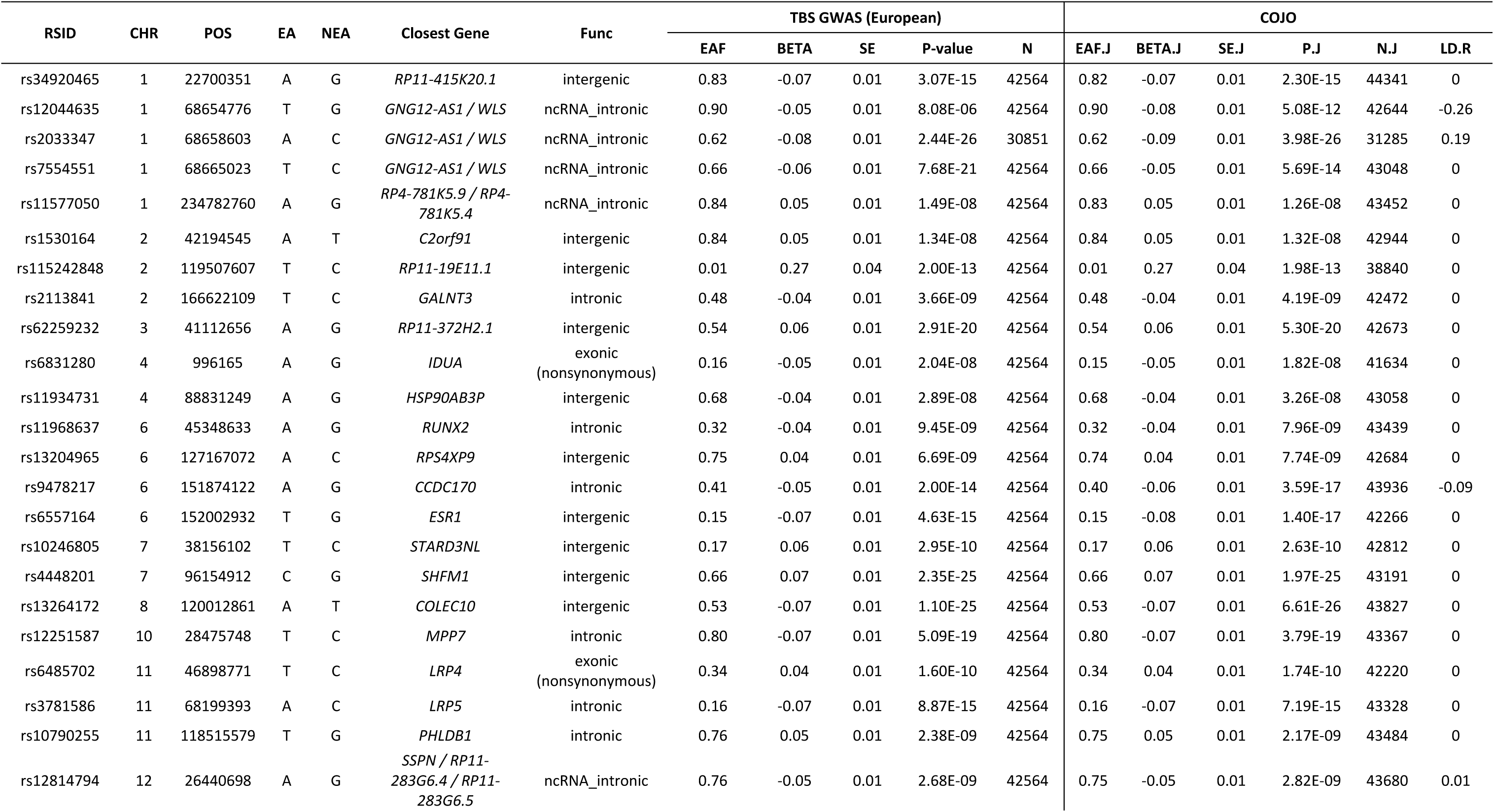
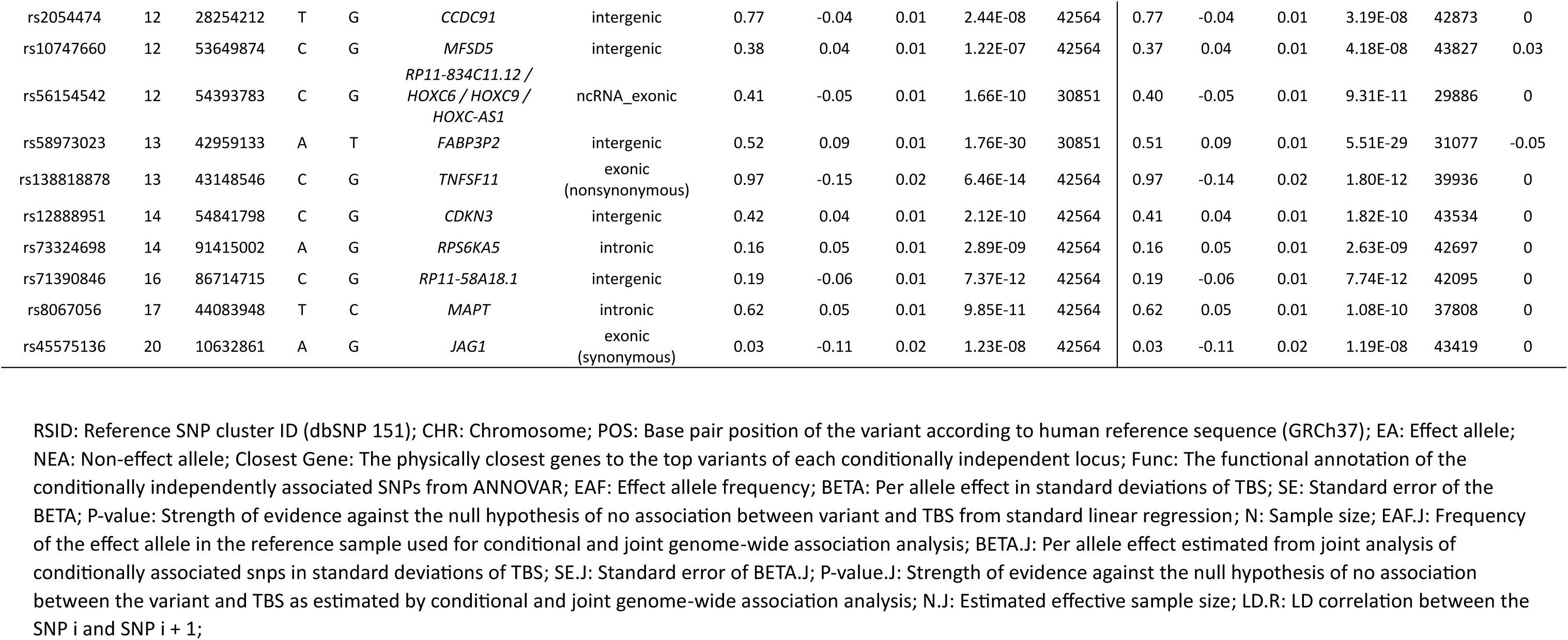
Conditional and joint (COJO) analysis identified 33 conditionally independent loci associated with TBS (P-value < 5E-8) from the European ancestry (N = 42,564).

After conditioning our results on the e-BMD associated variants^20^, we identified two independent signals mapping to *GALNT3* and *MPP7* that remained significantly associated with TBS (Supplementary Table 4). Nonetheless, these two variants have been reported as associated with TB-BMD^2^ and LS-BMD^50^.

### Sex heterogeneity of association effects

In our sex-stratified GWAS, 14 and eight loci were GWS-associated with TBS in females and males, respectively (Supplementary Table 5, Supplementary Figure 2). Associations in the vicinity of *GALNT3* (rs2113841) and *RAB11FIP3* (rs8059519) are GWS-associated with TBS in males. Whereas eight loci were GWS associated with TBS in females: *RP11-415K20.1* (rs7537281), *RP11-19E11.1*(rs115242848), *SFRP4*(rs11763529), *MPP7* (rs11006915), *C11orf49* (rs10769239), *LRP5* (rs3781586), *CDKN3* (rs6572952), and *RP11-58A18.1* (rs71390846). After multiple testing corrections, only the signal in *RAB11FIP3* showed significant sex heterogeneity (P-value: 2.33 ×10^−4^). The rs8059519-A allele was associated with higher TBS in males (Beta: 0.10, P-value: 4.34 ×10^−8^), but not in females (Beta: 0.01, P-value: 0.41) (Figure 2 and Supplementary Table 5).

**Figure 2:**
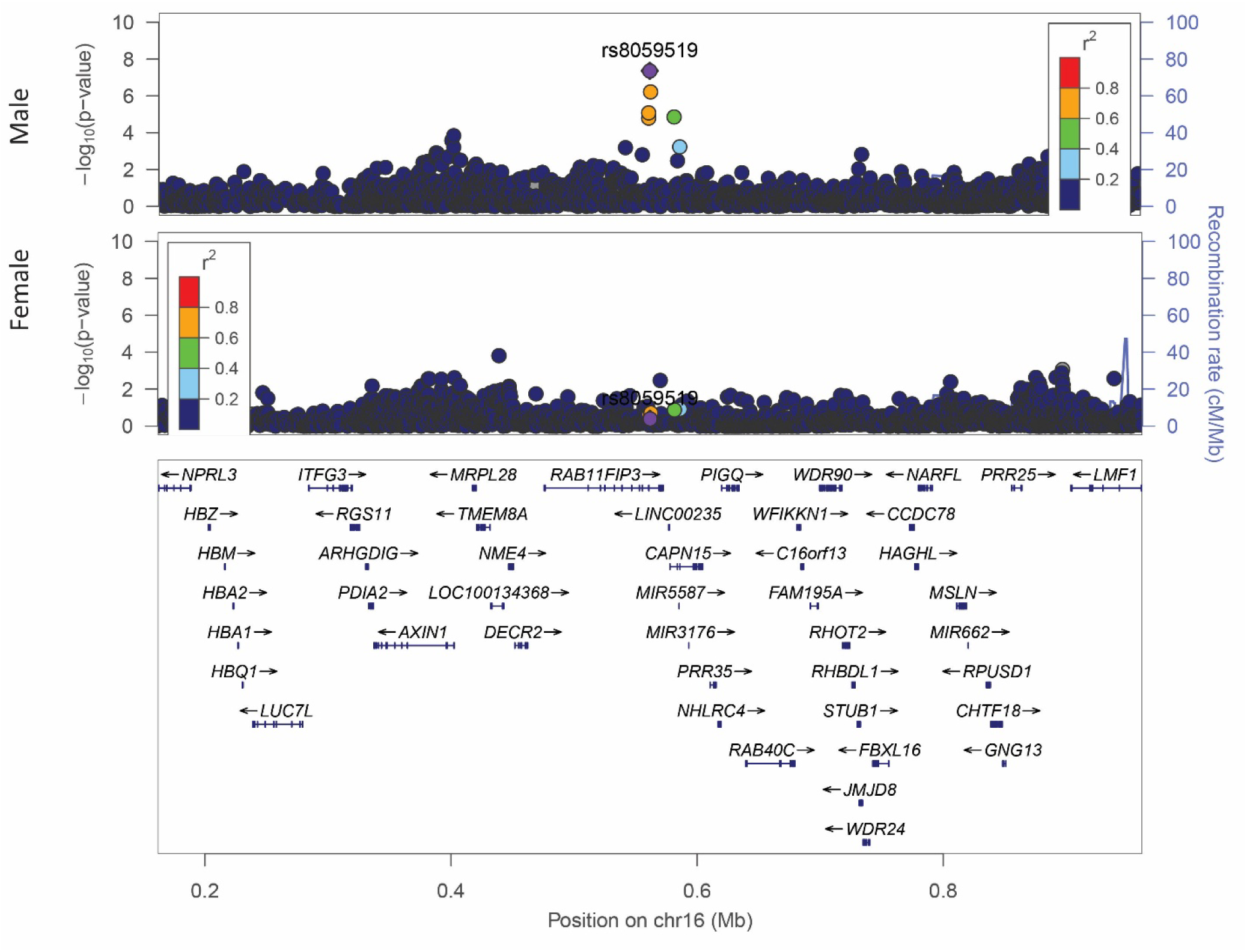
Regional plot (1,000kb) of rs8058519 from male- and female-specific TBS genome-wide association studies (GWAS). Regional plot showing the variant associations with TBS from male- (top) and female- (middle) specific GWAS and the protein-coding genes (bottom). The x- and y-axis indi ate the genomic position and –log10 transformed P-value of each SNP association, respectively. Colors of eac point indicate the linkage disequilibrium (LD) correlation coefficient (r2) of each variant with rs805951 (depicted in purple) from Locuszoom based on 1000 Genomes Project (European, Nov 2014). The blue lines i dicate the estimated recombination rates estimated from 1000 Genomes Project (European, Nov 2014).

### Functional annotation

From the 33 GWS TBS-associated variants in the European meta-analysis, five are located in exons, including rs6831280 (*IDUA*), rs6485702 (*LRP4*), rs138818878 (*TNFSF11*), rs45575136 (*JAG1*), and rs56154542 (*RP11-834C11.12:HOXC6:HOXC9:HOXC-AS1*) (Supplementary Table 6). Three of them, rs6485702, rs138818878, and rs56154542, are annotated as potentially deleterious with CADD scores of 19.99, 19.36, and 14.32, respectively. Moreover, rs6485702 was annotated as affecting the binding of transcription factors.

Genes mapping to TBS loci are significantly (FDR-adjusted P-value < 0.05) enriched in diverse biological pathways including skeletal system development, bone remodeling, skeletal system morphogenesis, clock-controlled autophagy in bone metabolism, type 1 collagen synthesis in the context of osteogenesis imperfecta, BMD and Alzheimer’s disease (Supplementary Table 7).

### Colocalization

Our results point to strong colocalization evidence (PWCoCo PP.H4 > 0.8) between TBS and cis-eQTLs (muscle skeletal tissue) in two regions: in 11p11.2 (rs6485702) with *ARHGAP1* expression (PP.H4: 0.97) and in 12q13.13 (rs56154542) with *HOXC6* expression (PP.H4: 0.95) (Supplementary Table 8). No evidence of colocalization was found between TBS-associated signals and gene expression measured in whole blood or osteoclasts.

### Genetic correlation

As shown in Table 2, Supplementary Figure 3, and Supplementary Table 9, TBS_sex-combined_ exhibits a positive genetic correlation with BMD measured at different skeletal sites (highest with LS-BMD, rg : 0.90, P-value: 3.81×10^−50^). Additionally, it also presents a negative genetic correlation with AnyFX (rg : −0.46, P-value: 5.96 ×10^−16^), HipFX (rg : −0.46, P-value: 1.23 ×10^−8^), and ForearmFX (rg : −0.42, P-value: 2.32 ×10^−12^), similar to the correlation pattern observed for LS-BMD. Similar patterns have also been observed from sex-stratified TBS GWAS (Table 2).

**Table 2:** The genetic correlation (rg) between TBS (sex-combined, male-only, and female-only) with skeletal traits, plausible risk factors, and diseases conducted using LDSC.

| Phenotypes | PMID | TBS (Sex-combined) |  |  | TBS (Male-only) |  |  | TBS (Female-only) |  |  |
| --- | --- | --- | --- | --- | --- | --- | --- | --- | --- | --- |
|  |  | rg | SE | P-value | rg | se | P-value | rg | se | P-value |
| estimated heel BMD | 30048462 | 0.50 | 0.04 | 1.64E-42 | 0.47 | 0.05 | 1.42E-20 | 0.52 | 0.04 | 4.18E-33 |
| Lumbar spine BMD | 26367794 | 0.90 | 0.06 | 3.81E-50 | 0.94 | 0.09 | 1.22E-24 | 0.88 | 0.07 | 1.51E-34 |
| Femoral neck BMD | 26367794 | 0.81 | 0.06 | 6.03E-43 | 0.86 | 0.09 | 9.91E-23 | 0.77 | 0.07 | 6.37E-31 |
| Skull BMD | 37402774 | 0.55 | 0.06 | 4.18E-22 | 0.55 | 0.07 | 1.47E-14 | 0.53 | 0.07 | 3.27E-15 |
| Total body BMD | 29304378 | 0.81 | 0.04 | 3.80E-102 | 0.80 | 0.06 | 5.60E-44 | 0.79 | 0.05 | 2.04E-50 |
| Any fracture | 30598549 | -0.46 | 0.06 | 5.96E-16 | -0.36 | 0.08 | 4.28E-06 | -0.52 | 0.07 | 2.74E-15 |
| Hip fracture | 36260985 | -0.46 | 0.08 | 1.26E-08 | -0.46 | 0.12 | 6.41E-05 | -0.41 | 0.09 | 5.08E-06 |
| Forearm fracture | 37919453 | -0.42 | 0.06 | 2.32E-12 | -0.45 | 0.07 | 5.73E-10 | -0.40 | 0.07 | 7.89E-09 |
| Hand grip | 29313844 | 0.07 | 0.04 | 0.11 | 0.09 | 0.05 | 0.11 | 0.06 | 0.05 | 0.22 |
| Whole body lean mass | 28724990 | 0.05 | 0.07 | 0.49 | -0.02 | 0.09 | 0.85 | 0.09 | 0.08 | 0.27 |
| Appendicular lean mass | 28724990 | 0.03 | 0.07 | 0.70 | -0.09 | 0.09 | 0.35 | 0.10 | 0.08 | 0.20 |
| Body height | 30124842 | 0.05 | 0.03 | 0.05 | 0.01 | 0.04 | 0.74 | 0.08 | 0.03 | 0.01 |
| Body mass index | 30124842 | 0.02 | 0.02 | 0.40 | -0.04 | 0.03 | 0.27 | 0.06 | 0.03 | 0.03 |
| Type 2 diabetes | 30054458 | 0.09 | 0.04 | 0.03 | 0.07 | 0.06 | 0.20 | 0.11 | 0.05 | 0.02 |
| 25-hydroxyvitamin D concentration | 32242144 | 0.04 | 0.03 | 0.13 | 0.06 | 0.04 | 0.15 | 0.04 | 0.04 | 0.30 |
| Coronary artery calcification | 37770635 | -0.09 | 0.06 | 0.12 | -0.04 | 0.08 | 0.67 | -0.14 | 0.07 | 0.06 |
| Coronary artery disease | 26343387 | -0.04 | 0.04 | 0.27 | -0.09 | 0.06 | 0.11 | -4.00E-04 | 0.05 | 0.99 |

Phenotypes: The skeletal traits, plausible risk factors, diseases; PMID: The unique reference number assigned to each publication indexed in the PubMed database; rg: Genetic correlation estimated from LDSC, expressed as the proportion of phenotypic variance that is shared due to genetic causes; SE: Standard error of rg; P-value: Strength of evidence against the null hypothesis of no shared phenotypic variance explained by common genetic variants;

Additionally, a nominal significant (P-value < 0.05) correlation was obtained between TBS (sex-combined and female-only) with T2D (rg_sex-combined_: 0.09, P-value_sex-combined_: 0.03; rg_female-only_: 0.11, P-value_female-only_: 0.02) and body height (rg_sex-combined_: 0.05, P-value_sex-combined_: 0.05; rg_female-only_: 0.08, P-value_female-only_: 0.01).

### Two-sample Mendelian randomization

Our results show that BMD and TBS are causally associated with fracture (Figure 3, Supplementary Table 10). Both a high genetically predicted FN-BMD (OR = 0.64, 95% CI: 0.60 to 0.69, P-value = 1.07 × 10^−32^) and TBS (OR = 0.63, 95% CI: 0.58 to 0.67, P-value = 7.54 × 10^−35^) are protective for AnyFX, ForearmFX (FN-BMD OR = 0.59, 95% CI: 0.53 to 0.65, P-value = 3.16 × 10^−25^; TBS OR = 0.66, 95% CI: 0.58 to 0.75, P-value = 3.23 × 10^−10^) and HipFX (FN-BMD OR = 0.52, 95% CI: 0.44 to 0.6, P-value = 7.72 × 10^−17^; TBS OR = 0.58, 95% CI: 0.49 to 0.68, P-value = 5.45 × 10^−11^). There is no evidence of violations to the MR assumptions from the sensitivity analyses (Supplementary Table 11).

**Figure 3:**
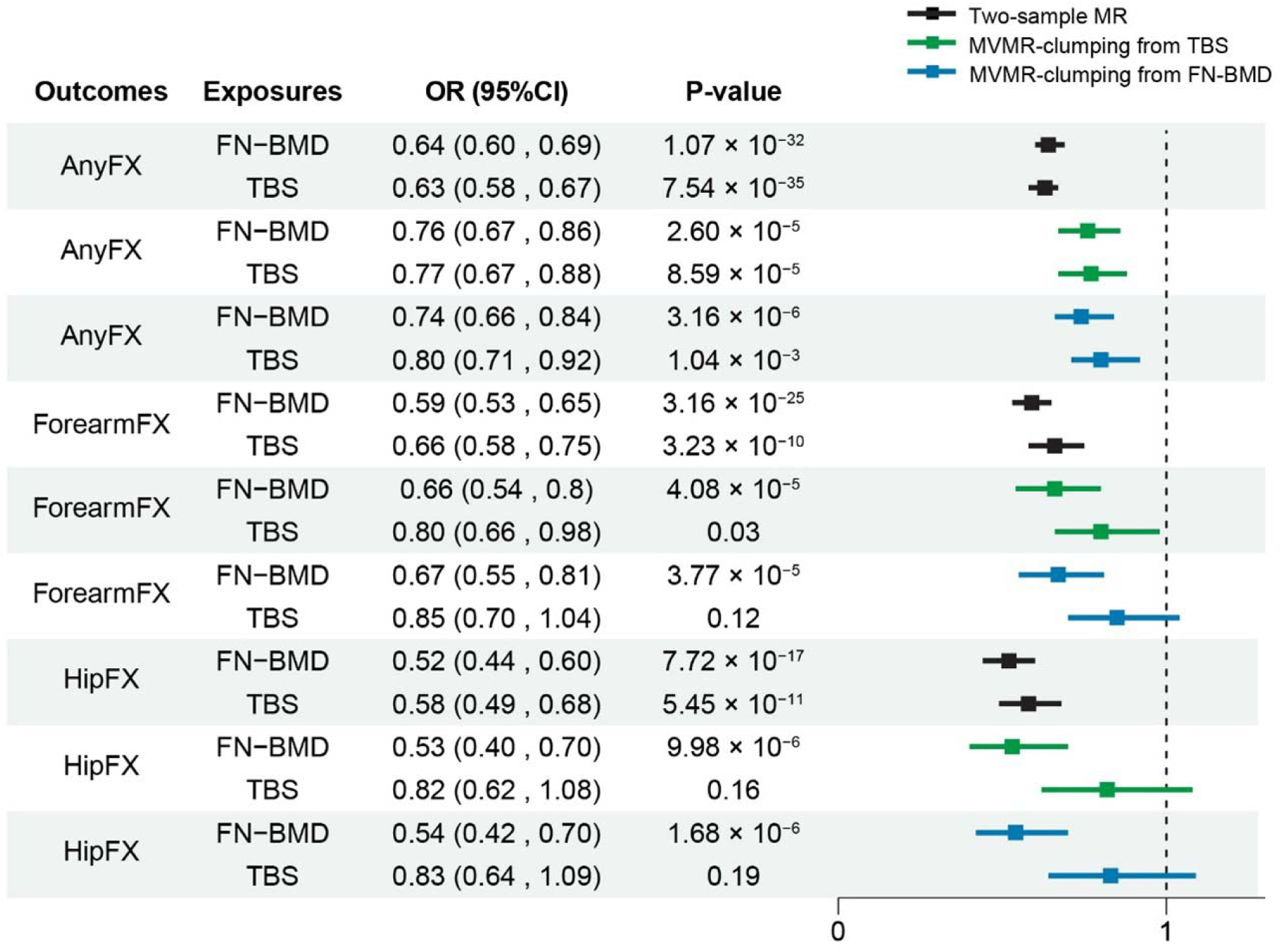
Forest plot indicates potential causal effects of FN-BMD and TBS on fracture risk based on two-sample Mendelian randomization (MR) and multivari ble MR (inverse-variance weighted) analyses. Forest plot illustrates the odds ratios (OR) and 95% confidence intervals for the estimated potential causal associations between femoral neck bone mineral density (FN-BMD), trabecular bone score (TBS), and fracture outcomes from different skeletal sites including any bone (AnyFX), forearm (ForearmFX), and hip (HipFX). The ORs are estimated using two-sample (black), MVMR () clumping based on variant associations with TBS (green), and MVMR clumping based on variant associations with FN-BMD (blue), respectively. The dashed vertical line represents the null effect (OR = 1).

### Multivariable Mendelian randomization

We extended the MR analyses to include FN-BMD (18 IVs) and TBS (26 IVs) as coexposures (Supplementary Table 12). Our MVMR (Table 3 and Figure 3) demonstrates a potentially causal association between TBS and AnyFX independent of FN-BMD, after clumping the IVs based on TBS summary statistics (OR: 0.77, 95%CI: 0.67 to 0.88, P-value: 8.59×10^−5^) or FN-BMD summary statistics (OR: 0.80, 95%CI: 0.71 to 0.92, P-value: 1.04×10^−3^). An independent effect of TBS on ForearmFX is observed using TBS summary statistics for clumping (OR: 0.80, 95%CI: 0.66 to 0.98, P-value: 0.03), but is rendered not significant when using FN-BMD summary statistics for clumping (OR: 0.85, 95%CI: 0.70 to 1.04, P-value: 0.12). Furthermore, there is evidence of weak instrument bias and significant heterogeneity effect for the coexposures in these analyses (i.e., conditional F-statistic less than 10) (Table 3).

**Table 3:**
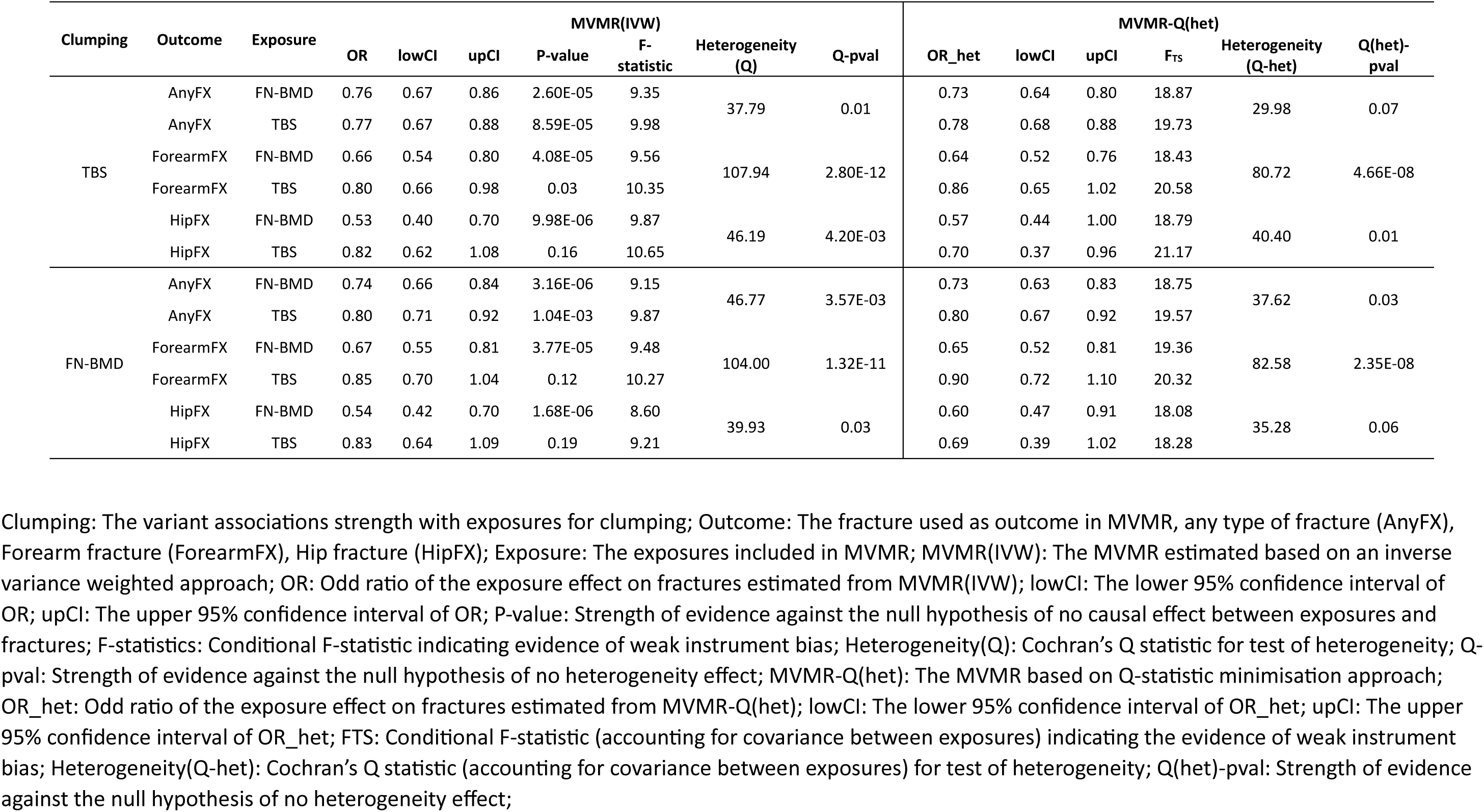
Multivariable Mendelian randomization (MVMR) indicating the direct causal effect of TBS on fractures at different bone sites that are independent of FN-BMD.

Accounting for the covariance between exposures using the MVMR-Q(het) provides a robust estimation of the coexposure effects even in the scenario of weak instruments and heterogeneity^48^ (Table 3). Using this method, we confirmed the independent causal effect of TBS on AnyFX when clumping variants based on the TBS summary statistics (MVMR-Qhet OR: 0.78, 95%CI: 0.68 to 0.88) or FN-BMD summary statistics (MVMR-Qhet OR: 0.80, 95%CI: 0.67 to 0.92). The results were less robust for HipFX, clumping based on TBS summary statistics resulted in an OR: 0.70, 95%CI: 0.37 to 0.96, whereas clumping based on FN-BMD deemed the association not significant (OR: 0.69, 95%CI: 0.39 to 1.02). No evidence of a BMD-independent effect was observed on ForearmFX.

Furthermore, we also used e-BMD (Supplementary Tables 13) as a coexposure in the TBS association with fracture, as it is the largest BMD study to date. As shown in Supplementary Table 14, with less-likely evidence of weak-instrument effect (conditional F-statistic > 10), TBS presents an independent causal relationship on HipFX (MVMR-IVW OR: 0.68, 95% CI: 0.54 to 0.84, P-value: 4.40×10^−4^). Using MVMR-Qhet, the effect estimate reduces in size but remains significant (OR: 0.59, 95% CI: 0.44 to 0.89).

Similarly, TBS presents a BMD-independent causal effect on AnyFX (MVMR-IVW OR: 0.83, 95% CI: 0.75 to 0.91, P-value: 2.04×10^−4^, MVMR-Qhet OR: 0.82, 95% CI: 0.67 to 0.99). No causal effect is observed between TBS and ForearmFX. The significant heterogeneity detected MVMR-Q(het) suggests this method provides more robust causal estimation as compared to MVMR-IVW (Supplementary Table 14).

## Discussion

In this study, we conducted the largest TBS GWAS to date, incorporating data of 44,767 participants from eight cohorts. Our analysis revealed 33 conditionally independent signals mapping to 29 loci associated with TBS. Altogether, these signals explain 4.06% of TBS phenotypic variance. In addition, we identified one locus (*RAB11FIP3*) exhibiting significant sex heterogeneity, with TBS-association observed only in males. All genomic regions identified as associated with TBS have previously been reported in association with BMD. Moreover, our MR results indicate that low TBS increases the risk of fractures, which appears to be independent of BMD and is consistent with evidence from observational studies that TBS could predict fractures over BMD^51^.

Within the GWS variants associated with TBS, we find nonsynonymous variants such as rs6485702, located in the exonic region of *LRP4*, with the T allele associated to increased TBS. Evidence from GTEx suggests this allele is significantly associated (P-value < 0.05) with decreased *LRP4* expression in the pancreas, liver, thyroid, and fibroblasts. LRP4 is a transmembrane receptor of sclerostin, which inhibits the Wnt/β-catenin signaling and bone formation^52^. Besides, it could also promote RANKL-induced osteoclast formation. An elevated cortical and trabecular bone mass has been observed in mice with *LRP4* loss in osteoblasts^52^. Additionally, rs6485702-T also decreases the expression of *ARHGAP1* in musculoskeletal tissue (P-value: 4.30 × 10^−23^), and this signal colocalizes with TBS (PP.H4: 0.97). ARHGAP1 is a negative regulator of mesenchymal stem cells (MSCs) osteogenic differentiation^53^. Consistent with our results, the rs6485702-T has also been reported to be associated with increased eBMD^1,20^ and TBBMD^2^.

Our study also shows that the C allele of rs138818878, a nonsynonymous variant in *TNFSF11*, is associated with decreased TBS. Plasma proteomics analyses from UKBB have shown that rs138818878-C is positively associated with TNFSF11 levels^54^. The TNFSF11 is an essential mediator of osteoclast formation, activation, survival, and could promote bone resorption^55^. TNFSF11 knockout mice present high bone mass due to the inhibition of bone resorption^56^. Similarly, rs138818878-C is associated with eBMD and skull BMD^1,3^.

We also found a signal mapping to *RAB11FIP3* (lead variant rs8059519-C), which is associated with TBS only in males. The protein family of Rab GTRase participates in cell trafficking processes^57^. The association of rs8059519-C with e-BMD has also been tested between males and females from e-BMD GWAS, however, no evidence of sex-heterogeneous effect was observed (P-value_het_: 0.72)^1^. Yet the potential mechanism of this locus that could affect trabecular bone differently in males and females should be further investigated.

Our Mendelian randomization analyses suggested there may be a causal effect of TBS on fracture risk. Although the results are robust to different sensitivity analyses, it is important to consider that there is a small (<7%) overlap between samples used to model the exposure (TBS) and the outcome (6.7% overlap with AnyFX GWAS^1^, 2.8% with forearmFX GWAS^30^ and 3.9% in the HipFX GWAS^29^), which could bias the MR results towards the observed association^58^. Given the low overlap rate, we do not expect this to impact our results significantly.

Assessing whether there is a TBS effect on fracture independent of BMD is rather difficult, given the high genetic correlation between the traits (e.g., rg between TBS and LS-BMD: 0.90). This high correlation is expected considering the same DXA image is used to derive both measurements. Besides, the genetic correlations between TBS and DXA-BMD (LS-BMD: 0.90, FN-BMD: 0.81) are higher than their phenotypic correlations (e.g., LS-BMD: 0.61, FN-BMD: 0.51, Pearson correlation calculated from 25,861 white British participants in UKBB), which is likely attributable to measurement error, hormonal, or environmental factors. To address the independence question, we have conducted MVMR, using FN-BMD as coexposure, as it holds a lower genetic correlation with TBS compared to LS-BMD and is not affected by the presence of osteophytes. Our results demonstrate a FN-BMD independent causal effect of TBS on hip and any-type of fracture. Nonetheless, FN-BMD was adjusted by sex, age, age squared, and weight in the original GWAS model^4^. And covariate adjustment in GWAS can bias the effect estimates obtained from MR studies^59^. Yet, using e-BMD as coexposure still detected an independent effect of TBS on fracture, and e-BMD was not adjusted for any heritable covariate^20^. An additional consideration is that our IVW analyses could have been biased by weak instruments. We have used the Q-minimization approach of MVMR-Q(het) and confirmed the causal association between TBS with any-type and hip fracture^48^. Nevertheless, to our knowledge, there are no examples in the current literature assessing the validity of MVMR in a scenario of such a high genetic correlation between exposures, and therefore our results should be interpreted with caution. On the other hand, triangulation offers credibility to the reported results as TBS has been shown to contribute to fracture risk prediction independently from BMD^10,51,60^.

The strength of our study lies in the large number of individuals included and the survey of sex-stratified effects. We also used genetic MR approaches to investigate the potential independent effect of TBS on fractures, by including large-scale GWAS of DXA and ultrasound heel BMD as coexposures, which are robust to potential observational confounders. However, some limitations need to be discussed. Our study predominantly focuses on individuals of European background, and although cohorts of other ethnicities are included in our analysis, they are of limited size. Therefore, future research is needed to elucidate whether our conclusions are also valid in other ethnicities.

In conclusion, this GWAS meta-analysis identified 33 conditionally independent variants associated with TBS at the genome-wide significance. We also report one locus with a sex-heterogeneity effect that is associated with TBS only in males. The MR analysis suggests that TBS may have a causal relationship with fracture risk, with a potential contribution beyond BMD; in line with observational studies evidencing the value of TBS on fracture risk prediction. Altogether, our study uncovers the genetic determinant of TBS and advocates for further investigation of bone traits other than BMD contributing to fracture risk.

## Contributors

Conceptualisation, Methodology: Haojie Lu, John P. Kemp, Carolina Medina-Gomez.

Data curation: Haojie Lu, John P. Kemp, Douglas P. Kiel, Griffin Tibbitts, Christine W. Lary, Monika Frysz, Gloria Hoi-Yee LI, Ching-Lung Cheung.

Formal analysis, Investigation: Haojie Lu

Writing—original draft: Haojie Lu, Carolina Medina-Gomez, John P. Kemp.

Writing—review and editing: all authors.

Supervision: John P. Kemp, Carolina Medina-Gomez.

Funding acquisition: Fernando Rivadeneira, Douglas P. Kiel, Nathalie van der Velde; Natasja van Schoor; Lissette de Groot; Andre G. Uitterlinden; Jonathan H. Tobias.

All authors read and approved the final version of the manuscript.

## Acknowledgments

**Rotterdam Study:** The generation and management of GWAS genotype data for the Rotterdam Study (RS-I, RS-II, RS-III) was executed by the Human Genotyping Facility of the Genetic Laboratory of the Department of Internal Medicine, Erasmus MC, Rotterdam. We thank Pascal Arp, Mila Jhamai, Marijn Verkerk, Lizbeth Herrera and Linda Broer, PhD, and Carolina Medina-Gomez, PhD, for their help in creating the GWAS database and for the creation and analysis of imputed data.

The Rotterdam Study is funded by Erasmus Medical Center and Erasmus University, Rotterdam, Netherlands Organization for the Health Research and Development (ZonMw), the Research Institute for Diseases in the Elderly (RIDE), the Ministry of Education, Culture and Science, the Ministry for Health, Welfare and Sports, the European Commission (DG XII), and the Municipality of Rotterdam. The authors are grateful to the study participants, the staff from the Rotterdam Study and the participating general practitioners and pharmacists.

Haojie Lu is sponsored by the PhD fellowship (202004910412) from the China Scholarship Council.

Fernando Rivadeneira is supported by LEGENDARE ERC-ADG 2020 101021500.

J. P.K. is funded by a National Health and Medical Research Council (Australia) Investigator grant (GNT2026272).

**Generation R**: We gratefully acknowledge the contribution of children and parents, general practitioners, hospitals, midwives and pharmacies in Rotterdam. The generation and management of GWAS genotype data for the Generation R Study was done at the Human Genomics Facility, HuGe-F, housed within the Laboratory for Population Genomics of the Department of Internal Medicine at Erasmus MC. We thank Zahra Alawi, Marijn Verkerk, Dr. Katerina Trajanoska, Costanza Vallerga, Samuel Gathan, Dr. Carolina Medina-Gomez, Dr. Linda Broer and Jard de Vries for the creation, management and QC of the GWAS database.

**Framingham Heart Study:** Research reported in this publication was supported by the National Institute of Arthritis and Musculoskeletal and Skin Diseases of the National Institutes of Health under award number R01 AR041398 and by Dairy Management Inc. The content is solely the responsibility of the authors and does not necessarily represent the official views of the National Institutes of Health. The Framingham Study was funded by the National Heart Lung and Blood Institute of the National Institutes of Health under contract N01-HC-25195, HHSN268201500001I.

Data is made available for this project through an approved data use agreement from dbGAP # 33361.

**UK Biobank:** This research has been conducted using the UK Biobank Resource (Application Number 17295), and was funded by a Wellcome Trust collaborative award (reference number 209233/Z/17/Z) which provided salary funding for MF. The phenotypic correlation analysis was conducted using the UK Biobank Resource under Application Number 67864. At the time this work was conducted MF was an employee at the University of Bristol. MF is now employed by Boehringer Ingelheim UK & Ireland.

## Data sharing

GWAS summary statistics of the Multi-ancestry, European ancestry, and sex-stratified meta-analysis will be made available through the GEFOS website with publication (http://www.gefos.org/).

