## Supplementary Figure for "The genetic architecture of trabecular bone score and association with fracture: a genome-wide association and meta-analysis"

### Supplementary Figures

**Supplementary Figure. 1: Miami plot of genome-wide association study (GWAS) of $TBS_{BMI}$ (adjusted by BMI) and $TBS_{thickness}$ (adjusted by tissue thickness).** This analysis was carried out from participants in UKBB (N = 28,534). On top is $TBS_{BMI}$ GWAS, and on the bottom is $TBS_{thickness}$ GWAS. The y-axis corresponds to the –log10 transformed P-value of each SNP association, and the x-axis corresponds to the SNP's base-pair position (GRCh37) along the chromosome. The dashed line indicates genome-wide significance (P-value < $5\times10^{-8}$).

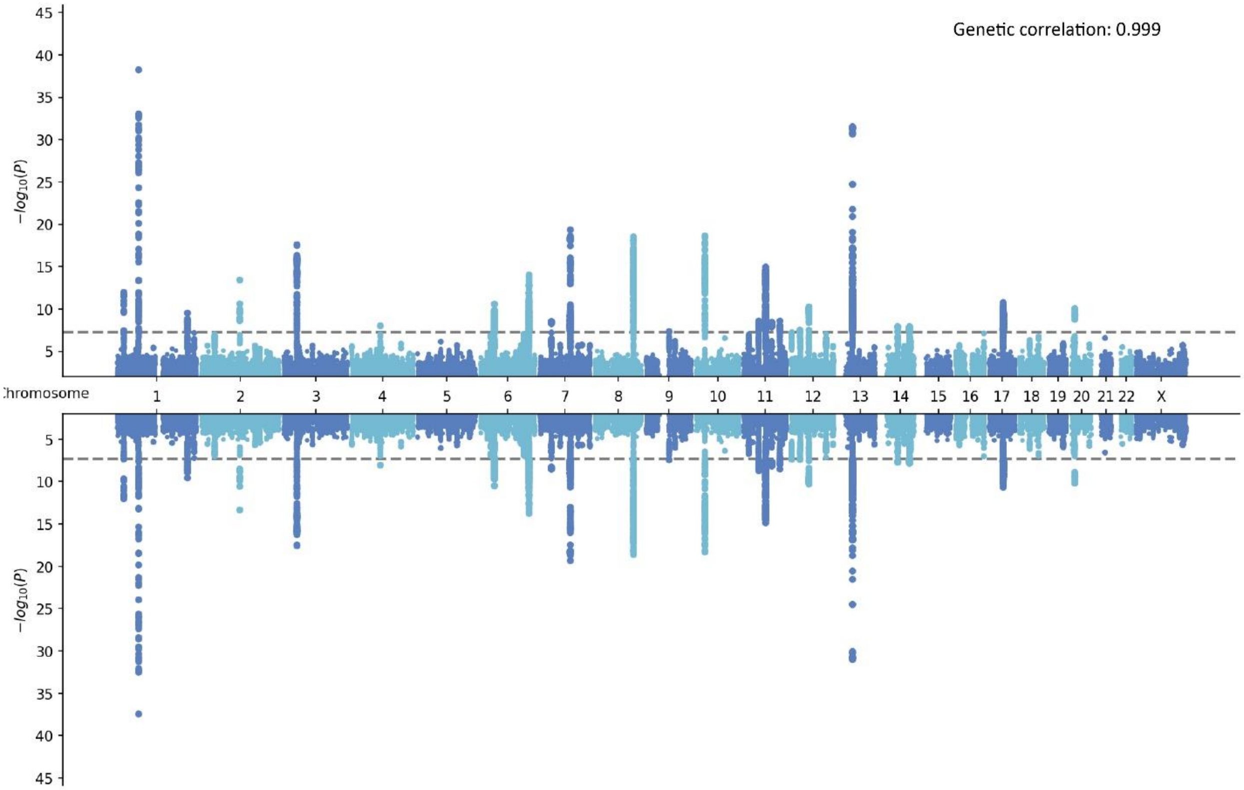

**Supplementary Figure. 2: The sex-stratified genome-wide association study (GWAS) of TBS**.

**A**: Quantile-quantile plot (left) and Manhattan plot (right) of TBS-GWAS for males only (N=19,118). **B:** Quantile-quantile plot (left) and Manhattan plot (right) of TBS-GWAS for females only (N=23,446). In Quantile-quantile plot (left), y-axis refers to the -log10 transformed P-value from TBS GWAS, x-axis represents the -log10 transformed expected P-value (two-sided). In the Manhattan plot, y-axis corresponds to the –log10 transformed P-value of each SNP association, the x-axis corresponds to SNP's base-pair position along the chromosome. The dashed line indicates genome-wide significance (P-value < $5\times10^{-8}$). The genes in the Manhattan plot refer to the closest gene to the lead variants of each locus.

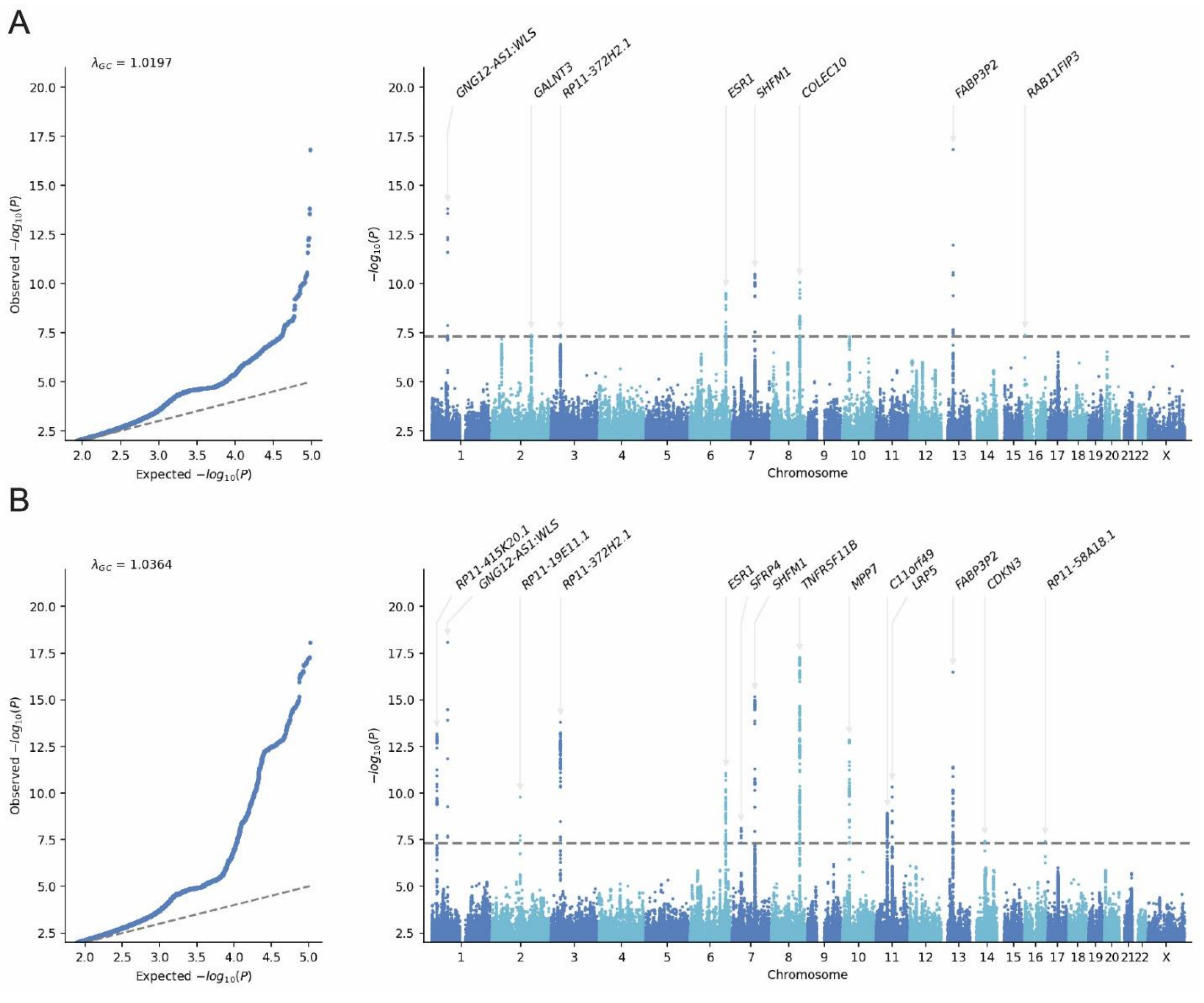

**Supplementary Figure. 3: Forest plot comparing the genetic correlation between TBS, LS-BMD, FN-BMD, and e-BMD with skeletal traits, plausible risk factors, and diseases conducted using LDSC.**

The dashed line indicates a genetic correlation of 0.

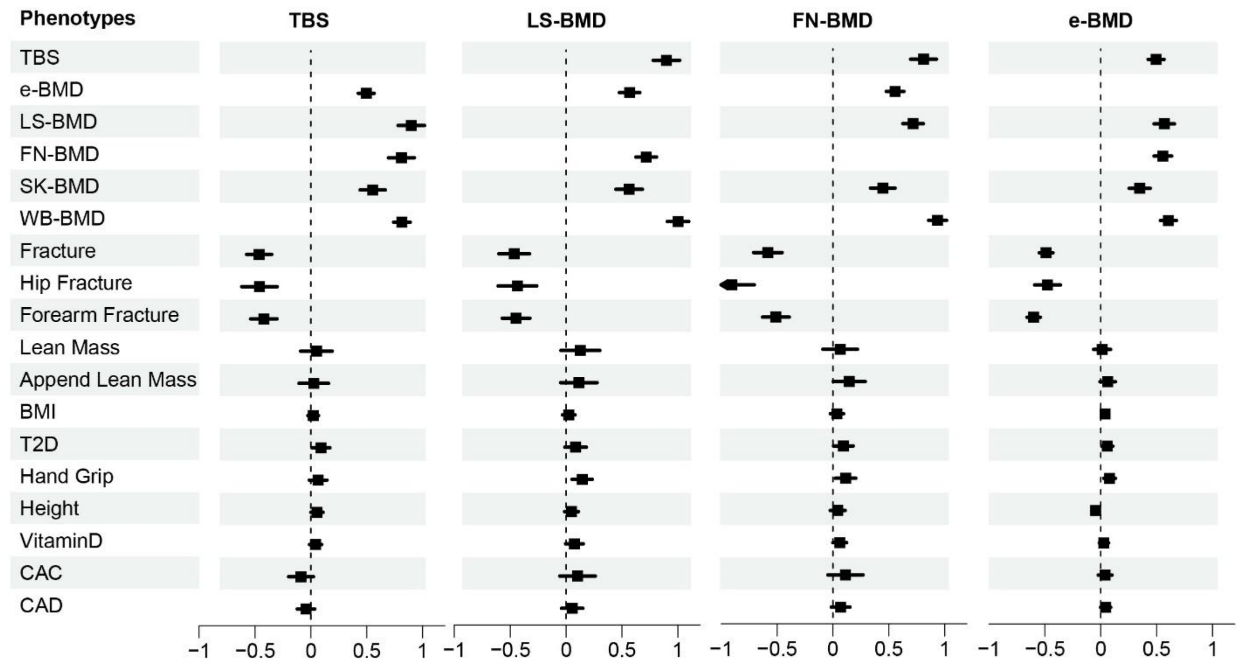
