## Supplementary Materials for "The genetic architecture of trabecular bone score and association with fracture: a genome-wide association and meta-analysis"

**Rotterdam Study:** The Rotterdam Study (RS) is a prospective population-based cohort study of older adults living in Ommoord, a suburb of Rotterdam, the Netherlands. Details of the study are described elsewhere[1]. It started in 1990 with the aim of describing the prevalence and incidence, unraveling the etiology, and identifying targets for prediction, prevention, or intervention of multifactorial diseases in mid-life and elderly. Currently this study includes 17,931 participants (overall response rate 65%), aged 40 years and over, who are examined in person every 3 to 5 years. Our study used participants from the fourth visit of RS-I, the third visit of RS-II, and the first visit of RS-III.

The Rotterdam Study has been approved by the Medical Ethics Committee of the Erasmus MC (registration number MEC 02.1015) and by the Dutch Ministry of Health, Welfare and Sport (Population Screening Act WBO, license number 1071272-159521-PG). The Rotterdam Study Personal Registration Data collection is filed with the Erasmus MC Data Protection Officer under registration number EMC1712001. The Rotterdam Study has been entered into the Dutch Trial Register (NTR; https://onderzoekmetmensen.nl) and into the WHO International Clinical Trials Registry Platform (ICTRP https://www.who.int/clinical-trials-registry-platform , search portal https://trialsearch.who.int/ ) under shared catalogue number NL6645 / NTR6831. The Rotterdam Study project persistant identifier is https://ror.org/02ac58f22.

All participants provided written informed consent to participate in the study and to have their information obtained from treating physicians.

**The Generation R Study**: The Generation R Study is a multiethnic prospective cohort study in which the general design, all research aims, and the specific measurements have been approved by the Medical Ethical Committee of Erasmus MC, University Medical Center Rotterdam. Details of study design and data collection can be found elsewhere[2]. There are totally 9,778 mothers with a delivery date from April 2002 until January 2006 were enrolled in Generation R. Measurements were planned in early pregnancy (gestational age <18 weeks), mid pregnancy (gestational age 18–25 weeks) and late pregnancy (gestational age >25 weeks).

Around the ages of 6 and 10 years, all children in Generation R and their parents were invited to participate in hands-on measurements, advanced imaging modalities, behavioural observations and biological sample collection in the Erasmus MC-Sophia Children's Hospital research center[2]. Our study used mothers at this visit with a gave written informed consent for each phase of the study.

**B-PROOF:** The B-Vitamins for the PRevention Of Osteoporotic Fractures study (B-PROOF) is a randomized double-blind placebo-controlled trial[3] which includes 2,919 participants aged 65 years and older with an elevated homocysteine concentration (≥ 12 µmol/L). The recruitment took place from August 2008 until March 2011. The B-PROOF study has been registered with the Netherlands Trial Register http://www.trialregister.nl under identifier NTR 1333 since June 1, 2008 and with ClinicalTrials.gov under identifier NCT00696514 since June 9, 2008. The WU Medical Ethics Committee approved the study protocol, and the Medical Ethics committees of EMC and VUmc gave approval for local feasibility[3].

All participants gave written informed consent before the start of the intervention.

**Framingham Heart Study:** The Framingham Osteoporosis Study (FOS) / Framingham Heart Study (FHS) is a family-based, multigenerational cohort Study initiated originally to study the risk factors for cardiovascular disease[4]. The FHS was initiated in 1948 to study determinants of cardiovascular disease and other major illnesses. The Original Cohort included 5,209 men and women, aged 28-62 years at enrolment who have undergone routine biennial examinations[5,6]. In 1971, Offspring of the Original Cohort participants and Offspring spouses including 5,124 men and women, aged 5 to 70 years, were enrolled into the Framingham Offspring Study. Offspring participants have been examined approximately every 4 years[7,8]. Framingham Heart Study investigators recruited the Third Generation Cohort (Gen3) from 2002 to 2005 (n=4,095 participants) (Am J Epidemiol 2007;165:1328–1335). In the 1990s, DNA was obtained for genetic studies from surviving Original Cohort Offspring participants. DNA from the Gen3 cohort was collected during the first two exams. This study included Offspring and Gen3 participants.

All studies were approved by IRBs from Boston University and the Advarra IRB for research conducted at Hebrew SeniorLife, and through approved data use agreement dbGAP #33361 at Northeastern University.

**UK Biobank:** In 2006-2010, the UK Biobank recruited 502,647 individuals aged between 37-76 years (99.5% were 40-69 years) from across the country[9]. Each participant provided information regarding their health and lifestyle using touch screen questionnaires, physical measurements and agreement to have their health followed and they also provided blood, urine, and saliva samples for future analysis. Data supporting this study was collected as part of the extended imaging study, which commenced in 2014[10].

UK Biobank has ethical approval from the Northwest Multi-centre Research Ethics Committee (11/NW/0382) and informed consent was obtained from all participants.

**HKOS:** The HKOS is a community-dwelling cohort comprising 9,449 Southern Chinese recruited from public roadshows and health fairs in various districts of Hong Kong in 1995-2010. Study participants were required to complete structured questionnaires, clinical and bone mineral density measurements at baseline. Since 2015, a full-scale in-person follow-up study commenced at which age-related parameters in addition to bone health were evaluated among the study participants[11].

Informed consent was obtained from the study participants and the study protocol was approved by the Institutional Review Board of the University of Hong Kong / Hospital Authority Hong Kong West Cluster (UW 03-140 T/140 and UW 15-236).

**Rerefence:**
